# Essential gene networks in acute myeloid leukemia identified using a *microRNA*-knockout CRISPR library screen

**DOI:** 10.1101/661140

**Authors:** Martino Gabra, Chiara Pastrello, Nicole Machado, Jonathan Tak-Sum Chow, Max Kotlyar, Tomas Tokar, Igor Jurisica, Leonardo Salmena

## Abstract

MicroRNAs (miRNA) are small RNAs that function as key modulators of gene expression. Due to their promiscuity of binding, a single miRNA may regulate several genes and hence, multiple pathways simultaneously. In addition, the 3’-UTR of mRNA can be recognized by several miRNA for suppression or degradation. We built a **mi**cro**R**NA-only **K**nock-**o**ut (miRKo) CRISPR/Cas-9 library to identify essential miRNA in Acute Myeloid Leukemia (AML) using OCI-AML2, OCI-AML3 and U937 cell lines as *in vitro* models. 10 miRNA were identified to be essential in our screen among all three cell lines: *miR-19b-1, -19b-2, 29b-2, -302a, -3678, -3713, -3910-1, -4447, -4718* and *-6795*. By using weighted degrees of association, we identified pathway hubs that uniquely affect all 3 cell lines by integrating miRNA:mRNA networks using mirDIP and pathway analysis using pathDIP. Through the miRKo screen, network membership analyses and biological anticorrelation scoring through patient data analysis, we identified *RRP2CA, RPS6KB-1, CREB1, RPM1A, MAPK10, MAP3K2, ITCH, FBX-W7, NR3C1* and *XIAP* as likely targets of the essential miRNA in AML; and signal transduction, apoptosis, TGF-beta signalling and MAPK signalling as candidate essential pathways in AML.

## Introduction

MicroRNAs (miRNA) are key post-transcriptional regulatory elements known to be conserved amongst plants, animals and some viruses. Formed by transcription from specific loci by RNA polymerases, they initially exist as <1 kb long transcript primary (pri)-miRNA. Within the nucleus they are prepared for export through splicing, capping, polyadenylation and packaging similarly to long stranded transcripts[1]. Further splicing and processing by DROSHA and PASHA transform the pri-miRNA into a pre-miRNA which can then be exported into the cytoplasm by EXPORTIN-5. When pre-miRNA is formed, it is folded into a secondary structure known as a “stem-loop”[1,2]. The stem-loop structure is subsequently recognized and processed by a cytoplasmic RNase III such as DICER into a dsRNA dimer which rapidly breaks down into two single strands, and depending on the stability of the single miRNA strand, either strand can be active[2–5]. A functional third miRNA formed from this complex is thought to originate from the loop region, known as loop-miRNA[6–7].

A mature miRNA is single-stranded ribonucleotide of approximately 19-25 bp in length which can guide argonaute (Ago) proteins and other essential components of the RNA-induced silencing complex (RISC) to target mRNAs for the transcriptional suppression or degradation[9,10]. miRNA recognition elements (MRE) on mRNA are commonly, but not limited to non-coding 3’-untranslated region (3’-UTR)[9–12]. Perfect complementarity between the miRNA and the mRNA is not required for full effects, and the stability of the miRNA is dependent on sequences both within and beyond seed region[3,13,14]. This non-stringent binding allows miRNA to regulate the expression of multiple RNA transcripts through target promiscuity and once bound, the endonuclease activity of the RISC is activated via the slicer activity of AGOI[1,13,15]. Multiple interactions between miRNA and mRNA form the central basis of complex miRNA-mRNA regulatory networks which affect 60% of protein coding transcripts and countless non-coding genes. Further complexity arises as a result of a single mRNA transcript harbouring several MREs[16]. miRNA have been demonstrated to play critical roles by modifying or controlling most major pathways and thus their dysregulation can affect many disease states including cancer. Hallmarks including cell division, self-renewal, apoptosis, and DNA damage response among others can be regulated by miRNA[17–22].

While there have been notable advances in the understanding and therapy of several cancers, there is a dearth of advancement in understanding the molecular underpinnings of Acute Myeloid Leukemia (AML). Recent efforts to advance the field of AML research have focused on improved stratification of poor responding versus high responding individuals through identification of leukemic stem cell signatures and residual disease status[23,24]. Although new targeted therapy strategies including monoclonal antibodies and small molecule inhibitors are constantly emerging, many of these strategies have not proven more effective than the standard of care with the exception of the use of all-trans retinoic acid (ATRA) in acute promyelocytic leukemia (APL) which has become nearly curable in the majority of cases[25]. For most AML patients, little has changed in terms of treatment with the standard of care continuing to be dual cytarabine and anthracycline therapy; a therapy fundamentally unchanged for the last 30 years[26]. The long-standing presence of this strategy is owed to its effectiveness with a mean response rate up to 70% and a lack of superior strategies for most AML subtypes[27,28]. Notwithstanding, drug resistance is also a major therapeutic challenge in the treatment of AML. Failure of initial therapy can be observed in up to 40% of AML patients, and even when initial therapy is effective, up to 70% of patients eventually succumb to their disease due to aggressive relapse within 5 years[29–31]. Therapies that modulate the growth of AML by targeting essential pathways may prove to be beneficial.

In AML, there are several distinct subtype specific miRNA expression patterns that are modified by known key leukemogenic genes and pathways including FLT3, NPM1, Core Binding Factor (CBF), TP53 and Ras-MAPK signaling[32–35]. In addition, miRNA have been demonstrated to affect and to be effectors of these genes and pathways working in concert with oncogenes or tumor suppressors. Consequently, miRNA can be effectively used as prognostic markers in AML[33,34,36]. Previous works in the dysregulations of *de novo* AML have revealed that specific miRNA such as *miR-126* are capable of controlling cell cycle progression and that it may regulate distinct pathways in AML versus normal hematopoiesis[37]. In addition, increasing or decreasing miRNA expression in preclinical models has demonstrated that modulation of miRNA can lead to anti-leukemic effects[38–43].

The ramifications of these observations may become more relevant to current cancer research in light of recent drug market breakthroughs of RNAi therapy with Patisiran, a novel RNAi therapeutic, which has recently entered the market[44]. These advances which are hailed by major advances in RNAi delivery and safety merits a deeper in-depth analysis of miRNA-based therapies in AML. To date, the 5 year survival remains at 25% for all patients and previously promising targets have proven to be impossible to treat using conventional methods[45]. Thus, we propose that the perturbation of miRNA expression and the gene networks they control, are critical for AML growth and that there are specific miRNA which are essential to AML growth and maintenance.

To date, comprehensive studies examining the role of miRNA in AML by a methodical CRISPR/Cas9 screen are lacking. Herein, we created a gRNA library that targets miRNA only, known as the miRNA knockout (miRKo) library. The successful use of CRISPR-Cas system for genome editing has transformed the landscape of genetic research as it offers greater coverage and higher efficiency than previous generations of drug screens such as siRNA library screens[46–49]. The CRISPR-Cas9 system has been shown to effectively knock down the expression of mature miRNA by introducing destabilizing indels as little as +/-1 base pairs in the stem or seed regions of a pre-miRNA[50]. CRISPR screens have been widely applied to identify genetic vulnerabilities of cancer and recent reports have demonstrated their power and applicability to human disease[51–53]. While specificity of gene targeting may improve in the coming years, miRNA screens have been made possible due to specific targeting of miRNA by gRNA within miRNA families[50]. We demonstrate the elucidation of key pathways is possible using this technology and that the results reflect survival data within clinical datasets.

## Results

### Generation of a miRNA-only CRISPR library and work-flow

We sought to study miRNA biology through high-throughput screens, however specific tools to do so are limited. To examine the functions of miRNA in a comprehensive manner, we have generated a <u>miR</u>NA-only <u>K</u>nock<u>o</u>ut (miRKo) CRISPR library. miRNA have an average length of 80bp and as such, the number of binding sites and the location of the binding site are limited **(Fig 1A).** The gRNA are chosen to be biased towards the 3’ or 5’ stem regions to disrupt proper miRNA folding and maturation (**Fig 1B**). They are also selected to have a slightly higher predicted binding score by the Azimuth 2.0 scoring method **(Fig 1C).** To generate the miRKo library, single guide RNAs (gRNAs) targeting all known miRNA-stem-loop sequences in the human genome were designed and an oligo library array was generated. The library was created such that each miRNA (1795 total miRNA) is targeted 3-4 times on average. The miRKo library comprises of 6,835 unique gRNAs that target 1795 miRNA 3-4 times. 1,000 non-targeting gRNAs (adapted from Gecko-v2) were also included in the array. (**Fig 1D**). Lentiviral vector compatibility was achieved by linking adaptor sequences by PCR to the gRNA oligo sequences and the gRNAs were cloned into the lentiGuide-puro backbone using Gibson cloning, validated, and packaged into virus (**Fig 1D**)[54]. Lentiviral miRKo particles were then titered and delivered to Cas9^+^ AML cell lines at a multiplicity of infection (MOI) of 0.3 at 300X representation. OCI-AML2, OCI-AML3 and U937 AML cell lines were chosen as they represent diverse AML classifications. After recovery, cells were maintained at 300X representation through passaging for 18 days. During this time, miRNA knockout which confers a proliferative advantage outcompeted non-targeting cells while miRNA knockouts which are essential for AML growth and maintenance will lead to cell death and a reduced representation of the respective gRNA (**Fig 1E**).

**Figure 1.**
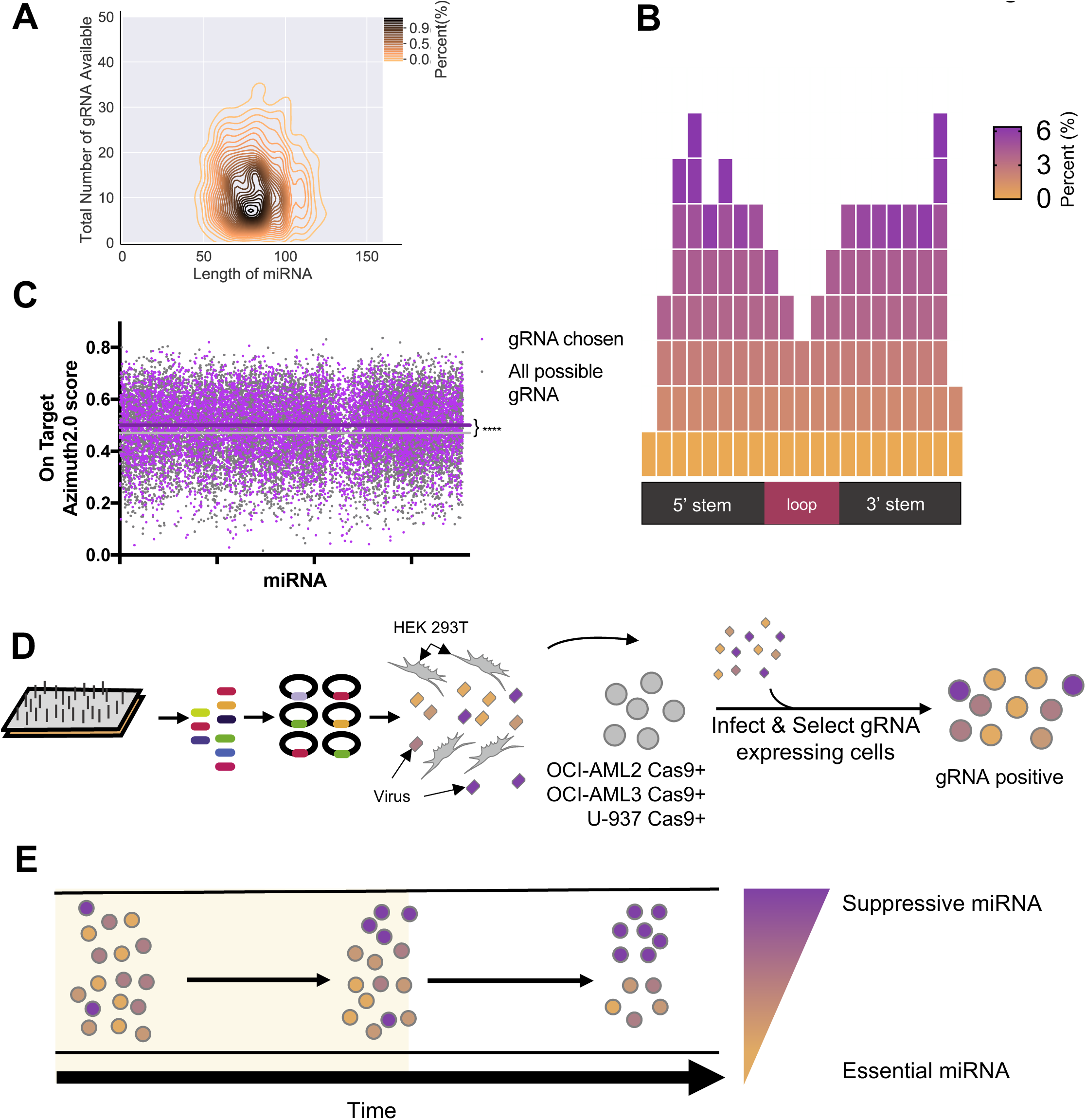
Creation of the microRNA knockout CRISPR library and project design. **A)** The average length of a miRNA transcript is around 80 nucleotides in length and contains a limited number of gRNA binding spots, with an average of 10 gRNA binding spots per miRNA. **B)** gRNA targeting miRNA are designed to be centered around the 5’ or the 3’ stem regions to disrupt the formation of the hairpin structure which leads to loss of mature miRNA formation.**C)** An average of 3-4 gRNA were selected per miRNA wherever possible and selection priority was given to gRNA with minimal off-target effect and higher on-target efficiency as determined by Azimuth 2.0 scoring. This resulted in a skew of higher on-target efficiency scores for selected miRNA (0.50±0.001,N=6835 vs 0.47±0.001,N=15057; p<0.0001) **D)** An array of oligonucleotides of all selected 6835 gRNA and 1000 non-targeting controls was created to be compatible with the lenti-guide puro plasmid (Addgene #52963) through a Custom Array generation and PCR amplification. Afterwards, lentivirus rich media was produced through the calcium/phosphate transfection of 293T cells and concentrated using LentiX solution. Parental AML cells (OCI-AML2, OCI-AML3 or U937) were infected with virus at an MOI of 0.3 at a representation of 300X, puromycin treated (1ug/mL) for 2 days and recovered for 3 days. **E) C**ells were then collected immediately for the initial time point and after growth for 3, 6, 9, 12, 15 and 18 days. gRNA changes are monitored for extreme changes where gRNA that increase growth reveal miRNA that suppress growth while gRNA loss represents miRNA that are essential and required for growth of cells.

### NGS Sequencing and Analysis

The gRNA within the three cell lines trended towards a net loss (**Fig 2A**). In addition, it was observed that targeting miRNA within all three cell lines had a similar effect on their growth such that miRNA which led to growth suppression in one cell line, trended towards growth suppression within the other cell lines as well. miRNA which were heavily suggested to be essential with a z-score greater than 3 were represented as essential within all 3 cell lines **(Fig 2B)**. Differential miRNA analysis identified 36 unique miRNA candidates at any time point until day 18, where 15 genes are found to be essential across all three cell lines **(Fig 2C).** By following the dropout status of each gRNA throughout the time course, we identify 10 miRNA that are significantly enriched throughout the time course from time point 6 until time point 18 consecutively, with 5 miRNA achieving significance later at time point 18 only **(Fig 2D).**

**Figure 2.**
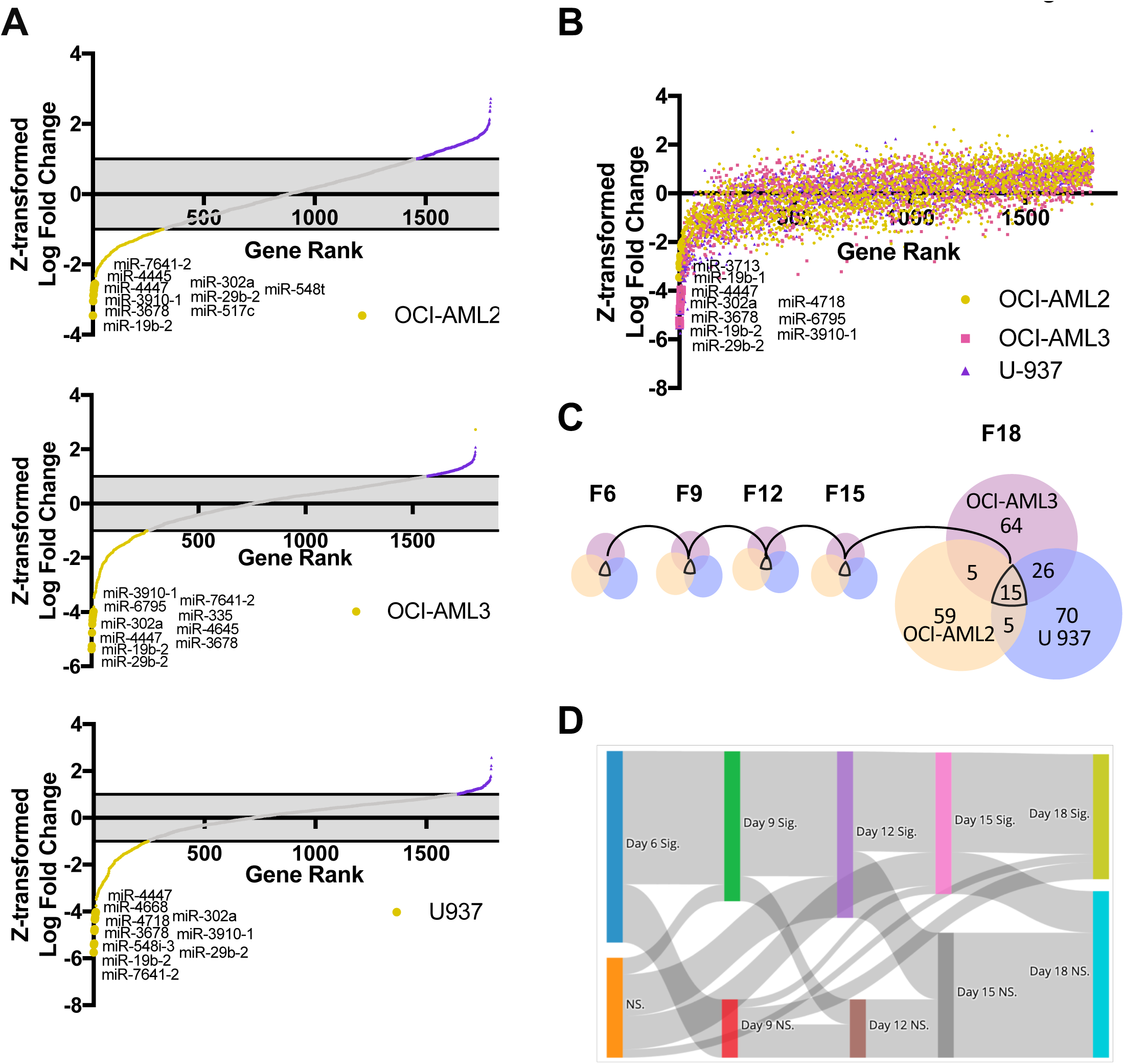
Identifying common targets affected within all OCI-AML2, OCI-AML3 AND U937. **A)** The z-transformed normalized score of the gRNA average fold changes at day 18 demonstrated that there is a trend to overall loss of gRNA and a maintenance of the miRNA with each cell line having unique essential miRNA as their top hits. **B)** The overall trend of changes at day 18 within the cell lines were reflected within each other suggesting that gRNA behaved in similar ways within each cell line. **D)** At any time point, there were 10-20 miRNA identified as commonly contributing to cell growth within all 3 cell lines. At time point 18, there were 15 miRNA that were identified as essential miRNA within all of the AML lines. **D)** By following the expression status of each gRNA set throughout the time course, it is observed that 10 of the 15 miRNA are identified as essential miRNA from time point 3 and onwards, and their respective gRNA are considered significantly lost throughout the time course.

Our miRKo screen resulted in the identification of 84, 110 and 116 candidate essential miRNA hits in OCI-AML2, OCI-AML3, U937, respectively **(Table S1)**. Among the essential miRNA identified, high confidence candidates include: *miR-19b-1, -19b-2, -29b-2, -302a, -3678, -3713, -3910-1, -4447, - 4718 and -6795*. These miRNAs represent the core essential miRNA observed in all time points within all 3 cell lines.

### Identification of the essential miRNA

The miRNA identified as essential genes demonstrated immediate loss of expression of the gRNA that was maintained until time point 18. For some of the identified miRNA, it was observed that gRNA which targeted the loop structure or the distal region of the stem structure were less likely to demonstrate a substantial fold change **(Fig 3A)**. For *miR-3910-1, -4447* and *-4718*, it is observed that gRNA that target the loop or very close to the loop structure often did not affect growth. gRNA that target the distal regions of miR-29b-2 and -302a also exhibit minimal effect. gRNA that exhibited maximal effect were centered near the mature miRNA sequence or at the stem regions; which would prevent formation of stable hairpin structures and prevent miRNA maturation. This screen identifies miR-19b-1 and miR-19b-2 as two core essential genes **(Fig 4A)**. These two precursor miRNA share a mature 3p region and a highly homologous 5p region however, the gRNA designed within this library are highly specific for their respective gRNA and the loss of either miRNA is independent of its homologous pair **(Fig 4B)**. In addition to identification within this screen, miR-19b-1 and miR-19b-2 are identified as having a growth promoting effect in AML lines alone when examined within the Genome CRISPR database **(Fig 4C)**. While both of these miRNAs exist within oncogenic clusters, the other miRNA within this cluster were not found as essential targets in this screen **(Fig 4D).** Furthermore, the expression of *miR-19b-1* and *miR-19b-2* correlated highly with each other with a pearson r value of 0.99 and they correlated highly with other miRNA such as *miR-19, -20a* and *-17*, but they did not correlate with *miR-363, -20b, -18b, -29a* and *-106a* **(Fig 4E)**. Within these two oncogenic clusters, miRNA that highly correlated with *miR-19b-1* or *miR-19b-2* were found to have a similarly high hazard ratio on overall survival in patients in the TCGA cohort with a correlation of 0.7 **(Fig 4F)**.

**Figure 3.**
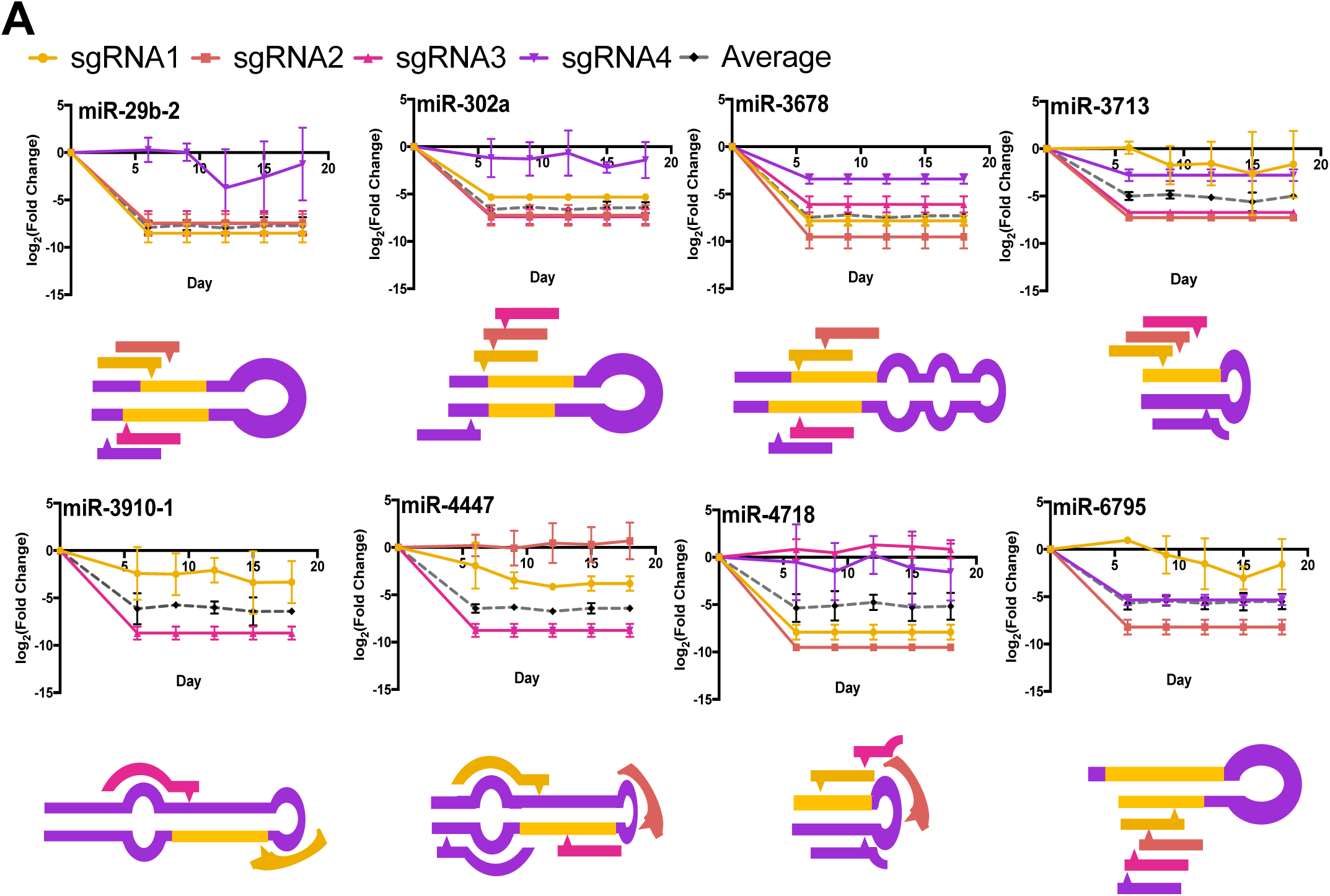
Profiling the Essential miRNA. **A)** Many of the identified miRNA demonstrate drastic effect of at least two gRNA and qualify as targets. Given the structure and availability limit of gRNA binding spaces on each miRNA, some gRNA target loop regions or distally on the stem structure. For miRNA such as miR-3910-1, -4447 and -4718, it is observed that gRNA that target the loop or very close to the loop structure elicit no effect on growth. gRNA that target the distal regions of miR-29b-2 and -302a also exhibit minimal effect. gRNA that exhibited maximal effect were often found near the mature miRNA sequence or at the stem regions; which would prevent formation of stable hairpin structures and prevent miRNA maturation.

**Figure 4.**
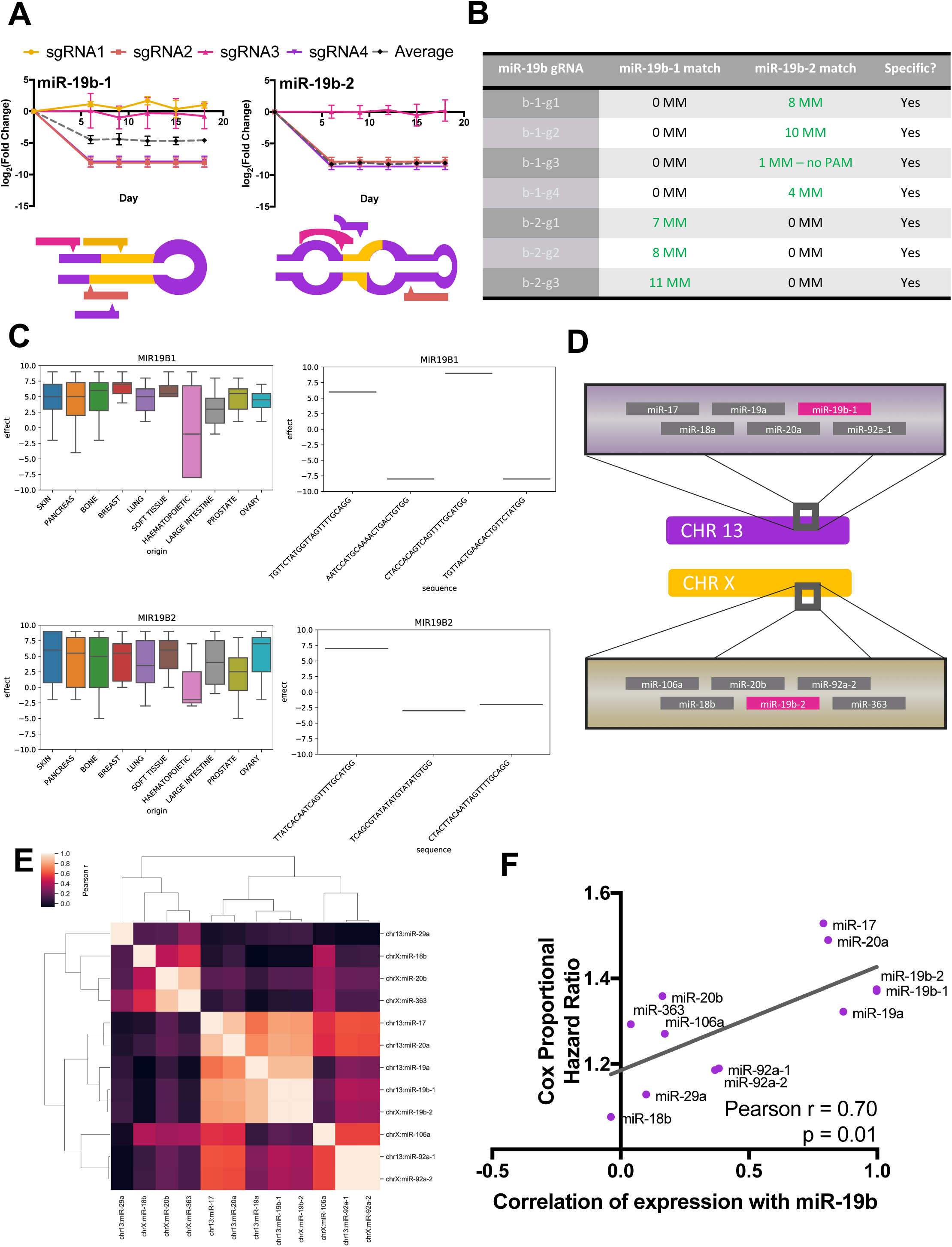
Identification of miR-19b-1 and miR-19b-2 as independent essential miRNA. **A)** Similarly to figure 3, miR-19b-1 and -19b-2 are identified as essential genes in OCI-AML2, OCI-AML3 and U937. **B)** Although the 3p sequence of miR-19b-1 and miR-19b-2 are identical and there is a high degree of overlap between the 5p fragments, the gRNA selected represent unique sequences that are specific for their respective transcript. This suggests that the identification of miR-19b-1 and miR-19b-2 as occurring independently of one another. **C)** By examining the GenomeCRISPR database for gene essentiality data, it is observed that both miR-19b-1 and miR-19b-2 are identified as potential essential miRNA. However, the study is limited to K562, a single AML cell line, and there is a discrepancy of effect between gRNA. Nonetheless, the gRNA targeting both miRNA show a trend towards essentiality within the leukemic cell line alone. **D)** The *miR-19b-1* and *miR-19b-2* transcripts represent miRNA derived from miRNA clusters from two different chromosomes: 13 and X. *miR-19b-1* is found within the *miR17HG* transcript, commonly known to be oncogenic, and containing *miR-17, -18a, -19a, 20a, - 19b-1* and *92a-1* while the second oncogenic cluster at the X chromosome contains the transcripts *miR-363, -92a-2, -19b-2, -20b, -18b* and *106a*. **E)** Hierarchical clustering of the expression of these transcripts demonstrates that despite the spatial separation between the two loci, there is a high degree of correlation (0.99) between the *miR-19b-1* and *miR-19b-2* transcripts. In addition, the expression status of these two miRNA closely follows the expression levels of some, but not all of the other transcripts found within these clusters. **F)** Within these loci, genes that were found to be highly correlated with *miR-19b* were found to similarly carry a high cox proportional hazard ratio (pearson r = 0.7).

### Profiling the downstream targets of the essential miRNA

miRNA are promiscuous in their target binding and therefore, multiple miRNA can converge on a single gene which may be implicated in a statistically enriched pathway **(Fig 5A)**. By using mirDIP, we generated computational likelihood scores of binding of miRNA to mRNA with scores of high confidence. Genes with high confidence of binding were assessed using TCGA expression data to generate anticorrelation values of miRNA-mRNA gene pairs. Interactions that were both anticorrelated and significant were retained in the pathDIP analysis **(Fig 5B).** Using the eigenvector centrality of each mRNA and the p-value generated of each pathway, we identify: RRP2CA, RPS6KB-1, CREB1, RPM1A, MAPK10, MAP3K2, ITCH, FBX-W7, NR3C1 and XIAP as the top 10 mRNA that participate within the essential miRNA network. These genes may exhibit their effect through pathways such as: signal transduction, apoptosis, TGF-beta signalling and MAPK signalling.

**Figure 5.**
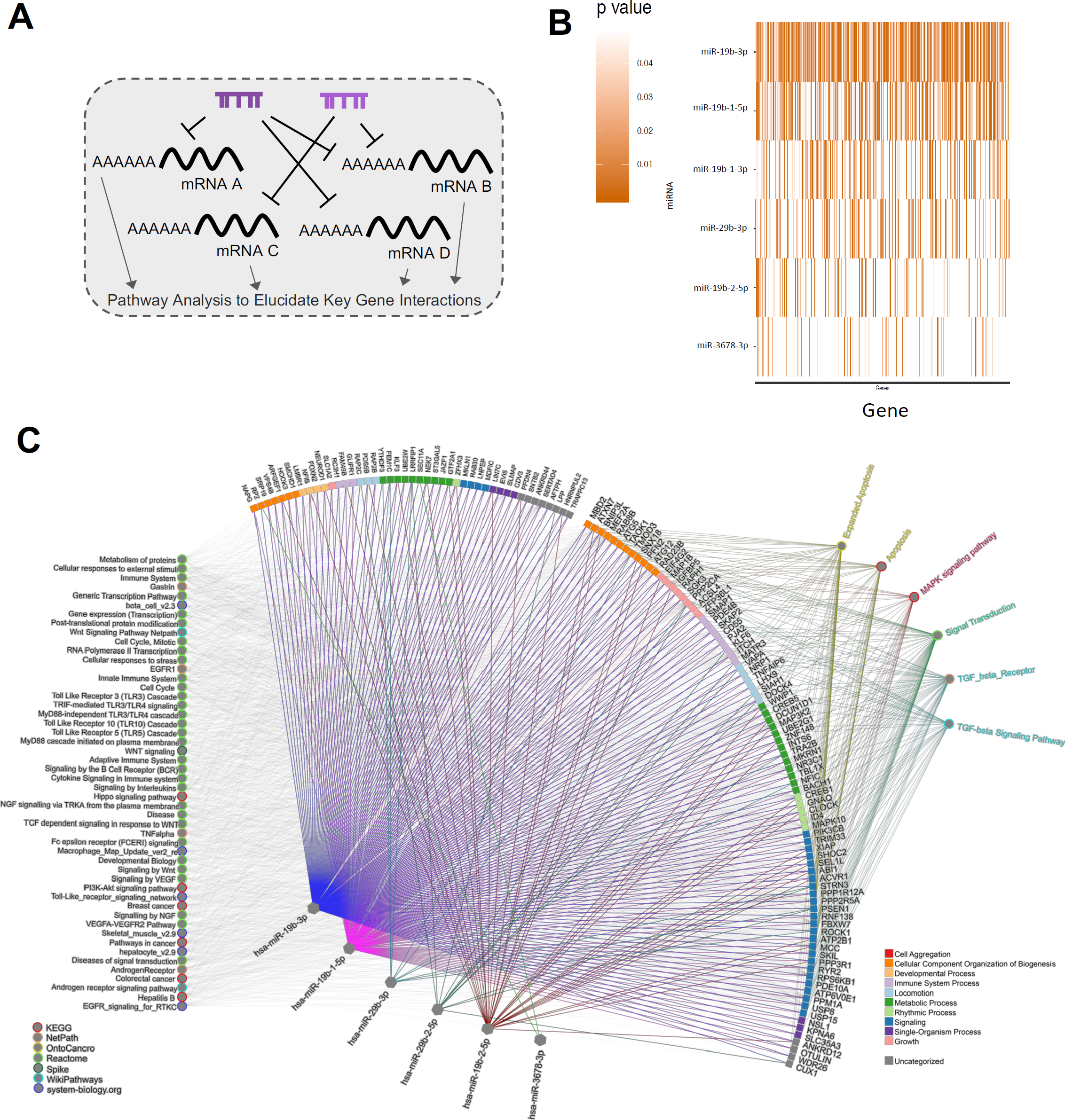
The pathway profile of the essential miRNA. **A)** Given that miRNA reduce the transcript level of mRNA, when a miRNA is lost due to a gRNA-induced mutation, the effect that the miRNA has on downstream pathways will be observed. When essential miRNA are lost, genes which inhibit proliferation are downregulated and the cell dies. Using mirDIP, we assess all possible mRNA-miRNA interactions and using pathDIP, we infer likely pathway interactions. **B)** We inform the computational targets discovered within mirDIP using TCGA data to assign correlation scores between the miRNA-mRNA interaction. Using the normalized expression data from RNA-seq and precursor expression data from the small RNAseq, we retain miRNA-mRNA gene interactions with a significant anti-correlation (rho<0, p<0.05). **C)** Using the eigenvector centrality of the mRNA targeted and the enrichment scores of each pathway as loading factors, we observe RRP2CA, RPS6KB-1, CREB1, RPM1A, MAPK10, MAP3K2, ITCH, FBX-W7, NR3C1 and XIAP as the top mRNA candidates likely targeted downstream of the essential miRNA in AML. These genes are involved in a variety of pathways including Signal transduction, apoptosis, TGF-beta signalling and MAPK signalling.

## Discussion

miRNA are promiscuous genes with several downstream targets and as such, they may have contextualized effects that are dependent on an intricate RNA landscape. Using a specialized miRNA knockout library (miRKO), we demonstrate that miRNA essential to a specific subtype of cancer can be deduced. With advanced computational tools such that integrate all major known scoring algorithms, downstream targets may also be elucidated. Within this analysis, we demonstrated that by examining top miRNA hits from three different AML lines, we can use eigenvector centrality and hub construction to predict common downstream targets and in turn, common downstream pathways associated with AML survival[55–57].

Although not much is known about the majority of the essential miRNA identified within this screen, we identify *miR-19b-1* and *miR-19b-2*, two miRNA from well known oncogenic clusters. These two miRNA have also shown some evidence of essentiality previously in a screen of K562 cells by Aguirre et al.[58]. This effect may also be unique to leukemic cells as a comparison of CRISPR screens between 36 cell lines demonstrates a selective negative selection of K562 cells[59]. These two miRNA are highly related by their mature miRNA sequences, differing only by 4-5 bases on the 5p fragment and having perfectly identical 3p sequences. Although they are located within two different well-known oncogenic loci, we observe a high degree of correlation of their expression in patient data obtained from TCGA. By examining the rest of the miRNA found at these two clusters, we find that *miR-19b-1* and *miR-19b-2* correlate the best with the most prognostic miRNA, namely: *miR-17, miR-19a* and *miR-20a*. They also seem to be least correlated to the least prognostic miRNA such as *miR-18b* and *miR-29a* (pearson r = 0.7). This would seem to suggest that the expression of these miRNA from these loci are not necessarily proportional, but that there is an underlying mechanism that facilitates the concomitant expression of the most oncogenic miRNA. Beyond the miR-19b gene pair, miR-29b-2, but not miR-29b-1 was identified within our screen. To date miR-29b-1 is highly implicated to be a tumour suppressor however, little is known of miR-29b-2. Patient data analysis of both the TCGA-LAML and the TARGET datasets demonstrates that high miR-29b-2 leads to a worse overall survival at the 75th percentile (data not shown)[60,61]. The remaining miRNA such as *miR-302a, -3678, -3713, -3910-1, - 4447, -4718* and *-6795* represent novel miRNA that are not well described in the context of AML propagation and survival. Further experiments to discern the effect of the gRNA on these miRNA is required as these miRNA are not well expressed in patient samples.

By using online databases such as mirDIP and pathDIP to deduce downstream targets and pathways, we discovered several common pathways of convergence for the essential miRNA. We predict which pathways are associated to coding genes highly enriched and anticorrelated to our miRNA of interest, indicating the value of affecting pathways such as: signal transduction, apoptosis, TGF-beta signalling and MAPK signalling.

Using the miRKo library and our experimental design, we identify miR-19b as uniquely important within the well-known oncogenic clusters miR17-92 and miR106a-363. We also identified several novel miRNA of unknown function as specifically enriched in our negative selection screen. It is possible that these genes may not be highly expressed or to have discreet effects at low expression status. Given the likelihood of interaction of each miRNA with several downstream gene targets, it is possible that signatures involving several miRNA may prove better predictors than any single miRNA alone[62].

In conclusion, the miRKo library demonstrates that miRNA dysregulations can be detected within each cell line. Using computational analysis, we demonstrate that gRNA targeting the core essential miRNA detected within all three cell lines are substantially downregulated. Of interest, *miR-19b-1* and *miR-19b-2*, two miRNA within key oncogenic clusters are independently identified as essential genes. Using computational and biological data, we demonstrate that these miRNA may act through common pathways such as: signal transduction, apoptosis, TGF-beta signalling and MAPK signalling. In future analyses, the miRKo library may also be used to identify pathways specific to drug resistance mechanism, clone propagation, leukemia initiation and other phenotypes where the role of miRNA is yet to be evaluated.

## Materials and Methods

### Suspension cell culture

OCI-AML2 and OCI-AML3 cells were maintained in alpha MEM media supplemented with 10% FBS and 1× P/S at 37°C and 5% CO_2_. U937 cells were maintained in RPMI with the same standard supplementation. All three suspension cell lines were maintained between 1.5×10^5 cells/mL and 1.0×10^6 cells/mL. 293T cells were maintained in DMEM media with standard supplementation.

### OCI-AML2-Cas-9, OCI-AML3-Cas-9 and U937-Cas-9 creation & validation

Virus rich media (VRM) of the Cas9-Blast plasmid (Addgene: #52962) was infected into regular AML lines using the standard lentivirus infection protocol described below. Blasticidin selection was conducted for 7 days at a dose of 1 ug/mL for all 3 cell lines. Afterwards, single colonies were propagated from the colony forming units (CFU) assay and Cas9 expression and function were validated by western blot to assess protein levels and a GFP-ablation assay by flow cytometry to assess Cas9 efficiency.

### Cas9 Activity Assay

Cells were assessed for Cas9 activity using the pXPR_011 plasmid (Addgene plasmid # 59702)[63] according to protocol by Doench *et al.* Briefly, cells were infected in a 6-well format at an MOI of 0.1-0.2 with the plasmid and 8 ug/mL protamine sulphate for 48 hours. After a 24 hour recovery period, cells were dosed with 2 ug/mL puromycin for 48 hours at a confluency of 1.5*10^5 cells/mL. Cells were allowed to rest for a minimum of 2 days prior to flow cytometry analysis using the Cytoflex. The pXPR_011 plasmid contains both a GFP sequence and a gRNA which cleaves this sequence. Consequently, the abrogation of fluorescence was used as a measurement of the efficiency of Cas9.

### Lentivirus production and infection

To create Virus Rich Media (VRM), 293T cells were plated at 1.5×106 cells confluency in 10 cm plates and they were transfected using calcium phosphate (2M CaCL2 and Hepes Buffered Saline) with 6.4 ug of psPAX2 (Addgene plasmid # 12260), 3.2 ug pMD2.G (Addgene plasmid # 12259) and 10 ug of the plasmid of interest. The media was replaced at 24 hours followed by collections at the 48 hour and 72 hour timepoints. Cell debris was removed by centrifugation at 500g for 10 minutes at 4 C and filtration with a 0.45 µm filter was conducted to ensure removal of cell debris.

To pellet the virus, we used a mixture of 26.4%w/v PEG 6000, 42 mM NaCl and 0.24X PBS at a ratio of 1 part mixture to 2 parts VRM. This mixture was placed at 4 C overnight on a rotating shaker and the virus was pelleted at 4500g for 45 minutes at 4 C afterwards. The resulting pellet was resuspended in PBS at a concentration of 20× and frozen at −80 C. To infect cells using the concentrated virus, we placed suspension cells at a concentration of 1.5×10^5 cells/mL, dosed the cells with the concentrated virus at an MOI of 0.3 (or below where specified) along with protamine sulfate (8 µg/mL) to improve infection. After 48 hours of incubation with the virus, cells were allowed to recover in regular media for 24 hours prior to blasticidin or puromycin selection (7 days or 2 days respectively).

### Determining the Multiplicity of Infection

To determine the multiplicity of infection (MOI), we conducted a dose finding analysis prior to each experiment. Briefly, cells were seeded at 1.5×10^5^ cells/mL in 2 mLs in 6-well dishes with varying volumes of virus and a constant dose of protamine sulfate (8 µg/mL). After infection and selection, viability of the cells were assessed to determine the concentration required for 30% cell survival.

### miRKo library infection

Three flasks of OCI-AML3, OCI-AML2 or U937 cells, clonally selected for Cas9 expression, were infected with the miRKo lentiviral library and treated with 8 ug/mL protamine sulfate to ensure a multiplicity of infection of 0.3. After 2 days of 2µg/mL puromycin (Bioshop) selection, the surviving cells from all of the plates were pooled together, mixed and split for 39 days, every 3 days.

### DNA Extraction for CRISPR insert sequencing

Genomic DNA was extracted from 3 × 10^6^ cells using Qiagen bloodamp mini kit (Cat No./ID: 51104). The resulting precipitate was treated with RNAse and pelleted using an ethanol prep and resuspended in EB buffer. DNA was extracted to ensure a 300× representation for miRKo infected cells at day 0 and day 39.

### miRKo library sequencing

To ensure appropriate gRNA representation, the input DNA for the first PCR had 300X coverage (∼16 ug of DNA). The conditions for the PCR largely followed the protocol used in Shalem *et al*.[64]. Briefly, an initial step to amplify the sequences off of the plasmid insert was conducted (18 cycles) followed by a step to add staggered barcode sequences through a second PCR (24 cycles). The primers for the first PCR are:

Fwd 5’ – AATGGACTATCATATGCTTACCGTAACTTGAAAGTATTTCG – 3’

Rev 5’ –CTTTAGTTTGTATGTCTGTTGCTATTATGTCTACTATTCTTTCC – 3’

While the second PCR primers were:

R2 5’ 5’–

CAAGCAGAAGACGGCATACGAGATGTGACTGGAGTTCAGACGTGTGCTCTTCCGATCTtctactattctttcccctgcactgt – 3’

F1 5’–

AATGATACGGCGACCACCGAGATCTACACTCTTTCCCTACACGACGCTCTTCCGATCTtAAGTAGAGtcttgtggaaaggacgaaacaccg – 3’.

F2

5’–

AATGATACGGCGACCACCGAGATCTACACTCTTTCCCTACACGACGCTCTTCCGATCTatACACGATCtcttgtggaaaggacgaaacaccg – 3’

F3

5’–

AATGATACGGCGACCACCGAGATCTACACTCTTTCCCTACACGACGCTCTTCCGATCTgatCGCGCGGTtcttgtggaaaggacgaaacaccg – 3’

F4

5’–

AATGATACGGCGACCACCGAGATCTACACTCTTTCCCTACACGACGCTCTTCCGATCTcgatCATGATCGtcttgtggaaaggacgaaacaccg – 3’

F5

5’–

AATGATACGGCGACCACCGAGATCTACACTCTTTCCCTACACGACGCTCTTCCGATCTtcgatCGTTACCAtcttgtggaaaggacgaaacaccg – 3’

F6

5’–

AATGATACGGCGACCACCGAGATCTACACTCTTTCCCTACACGACGCTCTTCCGATCTatcgatTCCTTGGTtcttgtggaaaggacgaaacaccg – 3’

### mirKo computational analysis

Fastq files were curated and trimmed to the same length using an original python script that was dependent on Biopython[65]. All sequences which lacked a barcode sequence within a prespecified region were discarded. After this trim, Bowtie2 was used to map reads to the library and to create a count table (<u>Langmead et al. 2009</u>). This count table was then used in Mageck-VISPR analyses for downstream analyses[66,67]. All p-values to identify essential genes are derived using this software, and data is visualized through python, R and graphPad Prism 6.

### TCGA data handling and mirDIP and pathDIP analyses

miRNA obtained from the screening were filtered based on their expression in parental cell lines (microRNAs not expressed in all parental cell lines were excluded) and in TCGA LAML samples (microRNAs not expressed in any TCGA sample were excluded). Precursor miRNA were then converted to mature using the miRBaseConverter package ver 1.6.0 (https://doi.org/10.1101/407148) in R 3.5.2. Prediction of targets for filtered microRNAs was conducted using mirDIP 4.1 with a setting of a minimum score of “High”[56]. TCGA mRNA and miRNA expression quantification were downloaded from https://portal.gdc.cancer.gov/ on 11/04/2019. Both files were log2-transformed and then normalized using the formula: □□□□□□□□=(□□□r-□□□□.*mol*)/□□.□□□.□□□ where mol can be either a miRNA or a gene, depending on the file.

Correlation between miRNA and gene expression was then calculated using Spearman correlation in R. Precursor miRNA-mRNA target pairs were then transformed to mature miRNA-mRNA target pairs, and only the pairs that were (i) present in our mirDIP prediction, (ii) had a rho<0 (anticorrelation), and (iii) had a p-value <0.05 were retained.

Anticorrelated targets were assessed for pathway enrichment using pathDIP using the “Literature curated” set[57]. The network of anticorrelated genes and microRNAs was used to calculate Eigenvector centrality using the igraph package (https://igraph.org/r/) in R.

TCGA data was also used to calculate Hazard Ratio for each microRNA and gene in our final list, using survival package (https://cran.r-project.org/web/packages/survival/index.html) in R.

## Acknowledgements

Leonardo Salmena is a recipient of a Tier II Canada Research Chair. This work was supported by funds to LS from a Career Development Award (CDA00079/2011-C) from the Human Frontier Science Program and an Operating Grant from the Leukemia and Lymphoma Society of Canada (LLSC; 569015). Martino Gabra is supported by a scholarship from the Centre for Pharmaceutical Oncology at the Leslie Dan Faculty of Pharmacy, University of Toronto and a trainee award from the LLSC. Computational analysis was in part supported by funds to IJ from Ontario Research Fund (#34876), Natural Sciences Research Council (NSERC #203475), Canada Foundation for Innovation (#225404, #30865), Canada Research Chair Program (CRC #225404), and IBM. pXPR_011 was a gift from John Doench & David Root (Addgene plasmid # 59702). psPAX2 was a gift from Didier Trono (Addgene plasmid # 12260). pMD2.G was a gift from Didier Trono (Addgene plasmid # 12259). lentiCas9-Blast was a gift from Feng Zhang (Addgene plasmid # 52962). Lentiguide-puro was a gift from Feng Zhang (Addgene plasmid # 52963).

---

**Supplementary Table S1.**
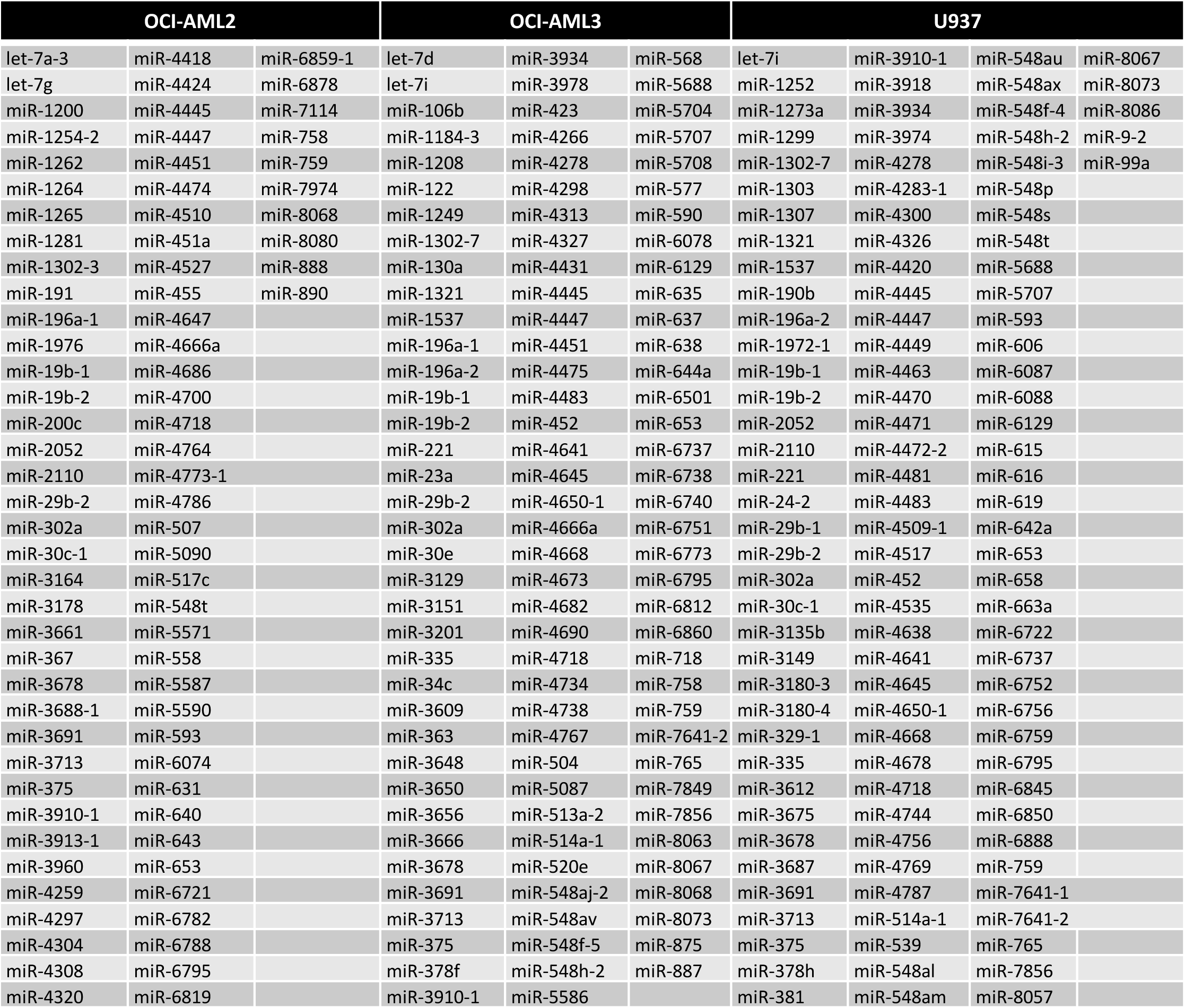
Essential miRNA.

