## Supplemental Table 1 for "Essential gene networks in acute myeloid leukemia identified using a *microRNA*-knockout CRISPR library screen"

### Supplementary Table S1. Essential miRNA

| OCI-AML2 | | | OCI-AML3 | | | U937 | | | |
|---|---|---|---|---|---|---|---|---|---|
| let-7a-3 | miR-4418 | miR-6859-1 | let-7d | miR-3934 | miR-568 | let-7i | miR-3910-1 | miR-548au | miR-8067 |
| let-7g | miR-4424 | miR-6878 | let-7i | miR-3978 | miR-5688 | miR-1252 | miR-3918 | miR-548ax | miR-8073 |
| miR-1200 | miR-4445 | miR-7114 | miR-106b | miR-423 | miR-5704 | miR-1273a | miR-3934 | miR-548f-4 | miR-8086 |
| miR-1254-2 | miR-4447 | miR-758 | miR-1184-3 | miR-4266 | miR-5707 | miR-1299 | miR-3974 | miR-548h-2 | miR-9-2 |
| miR-1262 | miR-4451 | miR-759 | miR-1208 | miR-4278 | miR-5708 | miR-1302-7 | miR-4278 | miR-548i-3 | miR-99a |
| miR-1264 | miR-4474 | miR-7974 | miR-122 | miR-4298 | miR-577 | miR-1303 | miR-4283-1 | miR-548p | |
| miR-1265 | miR-4510 | miR-8068 | miR-1249 | miR-4313 | miR-590 | miR-1307 | miR-4300 | miR-548s | |
| miR-1281 | miR-451a | miR-8080 | miR-1302-7 | miR-4327 | miR-6078 | miR-1321 | miR-4326 | miR-548t | |
| miR-1302-3 | miR-4527 | miR-888 | miR-130a | miR-4431 | miR-6129 | miR-1537 | miR-4420 | miR-5688 | |
| miR-191 | miR-455 | miR-890 | miR-1321 | miR-4445 | miR-635 | miR-190b | miR-4445 | miR-5707 | |
| miR-196a-1 | miR-4647 | | miR-1537 | miR-4447 | miR-637 | miR-196a-2 | miR-4447 | miR-593 | |
| miR-1976 | miR-4666a | | miR-196a-1 | miR-4451 | miR-638 | miR-1972-1 | miR-4449 | miR-606 | |
| miR-19b-1 | miR-4686 | | miR-196a-2 | miR-4475 | miR-644a | miR-19b-1 | miR-4463 | miR-6087 | |
| miR-19b-2 | miR-4700 | | miR-19b-1 | miR-4483 | miR-6501 | miR-19b-2 | miR-4470 | miR-6088 | |
| miR-200c | miR-4718 | | miR-19b-2 | miR-452 | miR-653 | miR-2052 | miR-4471 | miR-6129 | |
| miR-2052 | miR-4764 | | miR-221 | miR-4641 | miR-6737 | miR-2110 | miR-4472-2 | miR-615 | |
| miR-2110 | miR-4773-1 | | miR-23a | miR-4645 | miR-6738 | miR-221 | miR-4481 | miR-616 | |
| miR-29b-2 | miR-4786 | | miR-29b-2 | miR-4650-1 | miR-6740 | miR-24-2 | miR-4483 | miR-619 | |
| miR-302a | miR-507 | | miR-302a | miR-4666a | miR-6751 | miR-29b-1 | miR-4509-1 | miR-642a | |
| miR-30c-1 | miR-5090 | | miR-30e | miR-4668 | miR-6773 | miR-29b-2 | miR-4517 | miR-653 | |
| miR-3164 | miR-517c | | miR-3129 | miR-4673 | miR-6795 | miR-302a | miR-452 | miR-658 | |
| miR-3178 | miR-548t | | miR-3151 | miR-4682 | miR-6812 | miR-30c-1 | miR-4535 | miR-663a | |
| miR-3661 | miR-5571 | | miR-3201 | miR-4690 | miR-6860 | miR-3135b | miR-4638 | miR-6722 | |
| miR-367 | miR-558 | | miR-335 | miR-4718 | miR-718 | miR-3149 | miR-4641 | miR-6737 | |
| miR-3678 | miR-5587 | | miR-34c | miR-4734 | miR-758 | miR-3180-3 | miR-4645 | miR-6752 | |
| miR-3688-1 | miR-5590 | | miR-3609 | miR-4738 | miR-759 | miR-3180-4 | miR-4650-1 | miR-6756 | |
| miR-3691 | miR-593 | | miR-363 | miR-4767 | miR-7641-2 | miR-329-1 | miR-4668 | miR-6759 | |
| miR-3713 | miR-6074 | | miR-3648 | miR-504 | miR-765 | miR-335 | miR-4678 | miR-6795 | |
| miR-375 | miR-631 | | miR-3650 | miR-5087 | miR-7849 | miR-3612 | miR-4718 | miR-6845 | |
| miR-3910-1 | miR-640 | | miR-3656 | miR-513a-2 | miR-7856 | miR-3675 | miR-4744 | miR-6850 | |
| miR-3913-1 | miR-643 | | miR-3666 | miR-514a-1 | miR-8063 | miR-3678 | miR-4756 | miR-6888 | |
| miR-3960 | miR-653 | | miR-3678 | miR-520e | miR-8067 | miR-3687 | miR-4769 | miR-759 | |
| miR-4259 | miR-6721 | | miR-3691 | miR-548aj-2 | miR-8068 | miR-3691 | miR-4787 | miR-7641-1 | |
| miR-4297 | miR-6782 | | miR-3713 | miR-548av | miR-8073 | miR-3713 | miR-514a-1 | miR-7641-2 | |
| miR-4304 | miR-6788 | | miR-375 | miR-548f-5 | miR-875 | miR-375 | miR-539 | miR-765 | |
| miR-4308 | miR-6795 | | miR-378f | miR-548h-2 | miR-887 | miR-378h | miR-548al | miR-7856 | |
| miR-4320 | miR-6819 | | miR-3910-1 | miR-5586 | | miR-381 | miR-548am | miR-8057 | |
